# DeorphaNN: Virtual screening of GPCR peptide agonists using AlphaFold-predicted active-state complexes and deep learning embeddings

**DOI:** 10.1101/2025.03.19.644234

**Authors:** Larissa Ferguson, Sébastien Ouellet, Elke Vandewyer, Christopher Wang, Zaw Wunna, Tony K.Y. Lim, William R. Schafer, Isabel Beets

**Affiliations:** Neurobiology Division, MRC Laboratory of Molecular Biology, Cambridge, United Kingdom; Independent Researcher, Ottawa, Ontario, Canada; Department of Biology, KU Leuven, Leuven, Belgium; University College London, United Kingdom; University of Leeds, United Kingdom; Department of Pharmacology, University of Cambridge, Cambridge, United Kingdom

**Keywords:** G protein-coupled receptors (GPCRs), AlphaFold, deorphanization, peptide agonists, neuropeptides, peptide hormone, protein representations, active-state structures, protein embeddings, structural bioinformatics

## Abstract

Peptide-activated G protein-coupled receptors (GPCRs) regulate critical physiological processes such as metabolism, neural signalling, and endocrine function through their interaction with neuropeptides and peptide hormones. Despite their importance, identifying endogenous peptide agonists for GPCRs remains challenging, particularly for orphan receptors without known ligands. Recent advances in deep learning-based protein structure prediction, exemplified by AlphaFold (AF), have shown application beyond structural modelling, including protein-protein interaction prediction. Given that GPCR-peptide agonist interactions represent a specialized form of protein-protein interaction, we leveraged a dataset of experimentally validated agonist and non-agonist GPCR-peptide interactions from *Caenorhabditis elegans* to evaluate AF-Multimer’s ability to distinguish agonist-bound complexes. When modelling GPCR-peptide complexes, AF-Multimer confidence metrics partially discriminate agonist from non-agonist interactions, with improved discrimination achieved by utilizing AF-Multistate-derived active-state templates. We further investigated whether embeddings from the hidden layer of AF-Multimer’s neural network could distinguish agonist from non-agonist complexes. Feature performance analysis reveals that AF-Multimer’s pair representations outperform single representations, with distinct subregions of the pair representation providing complementary predictive signals. Building on these insights, we developed DeorphaNN, a graph neural network integrating active-state GPCR-peptide structural predictions, interatomic interactions, and deep learning embeddings to prioritize GPCR-peptide agonist interactions. DeorphaNN generalizes across datasets derived from different species, including annelids and humans, and successfully uncovered peptide agonists for two orphan GPCRs. DeorphaNN offers a novel computational resource to accelerate deorphanization by prioritizing GPCR-peptide agonist candidates for AI-guided experimental validation.

## INTRODUCTION

G protein-coupled receptors (GPCRs) are a large and diverse family of membrane proteins critical for cellular signalling, defined by their characteristic seven-transmembrane domain structure. A key class of GPCR agonists includes small peptides, such as neuropeptides and peptide hormones, which regulate physiological functions ranging from homeostasis and metabolism^1^ to complex behaviours like sleep, mechanosensation, and learning^2–5^. Peptide-activated GPCRs typically couple extracellular ligand binding to the regulation of intracellular signal transducers, mainly G proteins and arrestins^6^. However, identifying endogenous GPCR-peptide interactions remains a challenge, and many GPCRs are still “orphan” receptors without cognate agonists. Phylogenetic analyses have proven effective for predicting agonists of conserved peptide GPCRs across divergent phyla^7–11^, but are less effective for GPCRs without clear orthologues. Large-scale experimental screens that systematically evaluate extensive libraries of putative bioactive peptides derived through peptidomic analysis^12^ or computational tools^13–15^ have advanced peptide agonist discovery and GPCR deorphanization^9,11,15,16^, but these approaches are resource-intensive given that animal genomes typically encode dozens to over 100 peptide GPCRs alongside hundreds of bioactive peptides^9,15–17^.

Computational prediction methods can address these limitations by prioritizing high-probability GPCR agonist candidates for experimental validation. Given the therapeutic importance of GPCRs, numerous machine learning tools have been developed for *in silico* screening of small molecule GPCR ligands. For example, AiGPro uses a multi-task framework combining protein representations and SMILES strings to predict agonist and antagonist activities against human GPCRs^18^. DeepGPCR employs graph convolutional networks to classify small molecule-GPCR interactions^19^, while other models predict half-maximal effective concentration (EC_50_) values for chemical-GPCR pairs^20^. Structure-based docking tools, which model molecular interactions using physics-based simulations, have also proven effective for small molecule ligand discovery^21,22^.

Despite advances in machine learning applications and docking tools for identifying small molecule ligands for GPCRs, computational tools for *in silico* peptide agonist screening remain underdeveloped. This gap persists partly because peptides—larger, more flexible ligands than small molecules—pose unique challenges for computational modelling^23^. While docking methods can predict binding poses of peptide ligands of GPCRs^24^, they struggle to identify functional agonists^25^. This limitation highlights the need for next-generation computational tools specifically designed to address the unique structural and mechanistic complexities of GPCR-peptide agonist interactions.

Recent advances in protein structure prediction tools, including AlphaFold2 (AF2), are rapidly reshaping computational structure-based discovery of protein complexes and interactions. The AF2 algorithm^26^ predicts protein structure by implicitly learning biophysical relationships between residues through a combination of coevolutionary data and deep learning techniques^27^. These biophysical properties, encoded as statistical patterns in AF2’s protein representations, enable applications beyond structural modelling, including prediction of ligand-binding sites^28^, protein function^29^, and recombinant protein expression^30^. The development of AF-Multimer^31^, an extension of AF2, has allowed accurate prediction of multimeric complexes, including peptide-protein interactions^32,33^. Additionally, system-wide experimental screens are expanding our knowledge of GPCR-peptide interactions across diverse animal species, providing high-quality datasets for training machine learning models. In *Caenorhabditis elegans*, we recently constructed a comprehensive resource of over 400 GPCR-peptide agonist pairs and numerous non-agonist pairs through screening > 55,000 GPCR-peptide interactions^9^. Together, these advances—generalizable AF2-derived protein representations, AF-Multimer’s peptide-protein docking capabilities, and high-quality GPCR-peptide interaction data—provide a foundation for developing computational methods to predict peptide agonists for GPCRs.

In this study, we harness AF-Multimer-derived structural predictions and hidden layer protein representations to develop computational methods for classifying agonist and non-agonist GPCR-peptide interactions. Using a system-wide, experimentally validated *C. elegans* dataset of GPCR-peptide agonist and non-agonist pairs^9^, we evaluate AF-Multimer confidence metrics as indicators of agonist activity, revealing their modest discriminative capability. This capability is enhanced when active-state GPCR-peptide complexes— modelled using templates derived from AF-Multistate^34^—are utilized to align receptors with activation-associated binding conformations. Next, we analyze AF-Multimer’s hidden layer protein representations and find that pair representations outperform single representations in distinguishing agonists. We find that subregions within the pair representations contribute distinct, complementary predictive signals, highlighting the importance of multi-modal feature integration.

Building on these insights, we developed DeorphaNN, a graph neural network (GNN) that integrates AF-Multimer-derived structural predictions, interatomic interactions, and protein representations to classify agonist interactions. We demonstrate that this pipeline generalizes across datasets derived from different species, including *Platynereis* and human datasets. Experimental validation of top-ranked peptide agonists—prioritized by DeorphaNN—identifies endogenous peptide ligands of two orphan *C. elegans* GPCRs, demonstrating DeorphaNN’s potential for enhancing conventional screening and accelerating the deorphanization of peptide-activated GPCRs.

## RESULTS

### Dataset preprocessing

We leveraged a previously published dataset that employed a high-throughput CHO Gα_16_-mediated calcium signalling assay to screen peptide agonist activity across 161 putative peptide-activated GPCRs from *C. elegans*^9^. We only included GPCRs that showed concentration-dependent activation by at least one peptide, as a lack of activity could have been due to factors such as non-functional expression, misfolding, or failure to couple with Gα_16_, rather than the absence of a true agonist. Because AF-Multimer is unable to account for post-translational modifications, pyroglutamated and amidated residues were not included in peptide sequences. Following additional filtering—such as removal of short peptides (< 3 residues) and redundant GPCR and peptide sequences (see Methods)—our final dataset contained 65 GPCRs (derived from 55 genes) and 339 unique peptides (derived from 93 genes) (**Figure 1A**). Combining each GPCR with each peptide (“GPCR-peptide complex”) yielded 20,035 pairs, 457 of which were experimentally confirmed agonist matches^9^. The median number of agonists per GPCR in the dataset is 3 and ranges from 1-61 (**Figure 1B**). For 38 of the 65 GPCRs, agonist peptides are derived from a single peptide encoding gene, with the specific gene varying across receptors (**Supplemental Figure 1A**). Peptides range in size from 3 residues to 37 residues, with a median length of 11 residues (**Supplemental Figure 1B**). The names and primary amino acid sequences of all GPCRs and peptides used in this study are provided in **Supplemental Data 1**.

**Figure 1:**
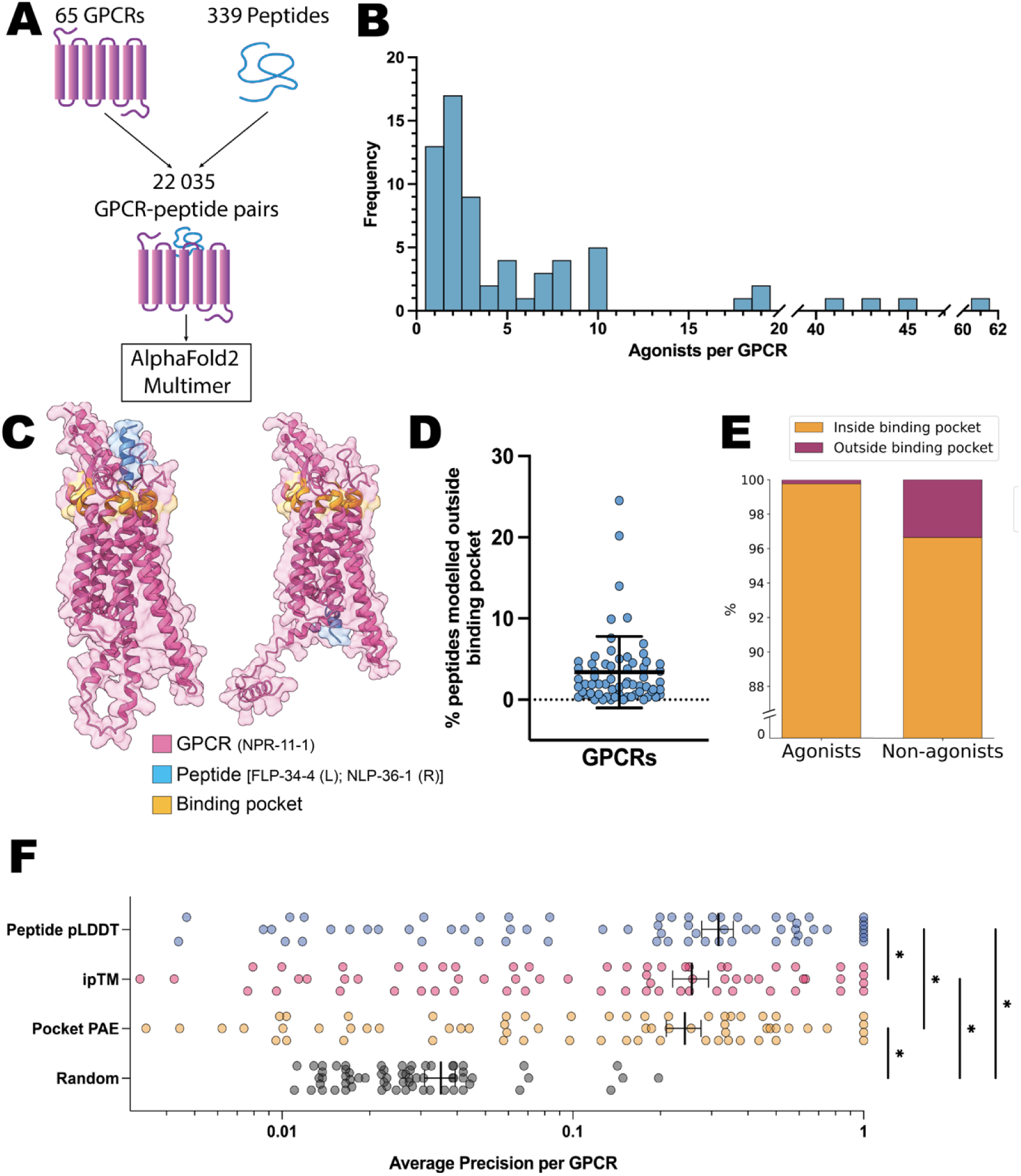
AF-Multimer structures and confidence metrics exhibit limited ability to discriminate between agonist and non-agonist GPCR-peptide interactions. (**A**) 65 GPCRs and 339 endogenous peptides (22 035 GPCR-peptide pairs) from *C. elegans* were modelled using AF2-Multimer. (**B**) Frequency distribution of the number of agonists per GPCR in the dataset. Most GPCRs have few agonists (median = 3), though a small number of promiscuous GPCRs account for the majority of agonist interactions. (**C**) Example AF-Multimer predicted structures. Left: FLP-34-4, an agonist for the GPCR, NPR-11-1, is modelling in the orthosteric binding pocket. Right: NLP-36-1, a non-agonist for NPR-11-1, is modelled outside of the binding pocket. The GPCR is shown in pink, binding pocket residues (defined as ±5 residues from the extracellular-membrane boundary) in yellow, and peptides in blue. (**D**) The percentage of peptides modelled outside the binding pocket for each GPCR ranges from 0 – 24.5%, with a mean of 3.4% (±4.4% SD).(**E**) 99.8% of agonists are modelled within the GPCR binding pocket, compared to 96.6% of non-agonists. (**F**) Peptide pLDDT significantly outperforms ipTM, pocket PAE, and random ranking in distinguishing agonists from non-agonists (Friedman test with multiple comparisons with FDR correction using the method of Benjamini, Krieger, and Yekutieli, * q < 0.05). Error bars indicate mean ± SEM.

### AF-Multimer positions peptides in GPCR binding pocket with minimal discrimination

We modelled each GPCR-peptide complex using AF-Multimer to evaluate whether the outputs could distinguish agonists from non-agonists. As experimentally determined structures show that peptide agonists typically interact with the helical cavity (“orthosteric binding pocket”) and extracellular loops of GPCRs^35^, we hypothesized that AF-Multimer would predominantly model agonist peptides in a similar manner, while non-agonist peptides might be modelled in less physiologically relevant locations (**Figure 1C**). To investigate this across diverse GPCR structures, we defined the putative GPCR binding pocket as residues within ±5 positions from the DeepTMHMM^36^-predicted extracellular-membrane boundary.

We then computed the minimum peptide-to-pocket residue distance to classify peptides as being “inside” or “outside” the GPCR binding pocket, using a 12.5 Å cutoff. On average, < 5% of peptides per GPCR were predicted to be further than 12.5 Å from the nearest binding pocket residue, though this varied by GPCR (**Figure 1D**). Across the dataset, only one agonist peptide was modelled outside of the binding pocket of its respective GPCR, while approximately 3.4% of non-agonist peptides were modelled outside the binding pocket (**Figure 1E**). Due to the prevalence of peptides positioned in the binding pocket irrespective of ligand class, this feature alone does not provide a reliable basis for distinguishing between agonist and non-agonist GPCR-peptide complexes.

### AF-Multimer confidence metrics partially discriminate peptide agonists from non-agonists

We next evaluated the ability of AF-Multimer confidence metrics (interface pTM [ipTM], predicted local distance difference test [pLDDT], and predicted aligned error [PAE]) to discriminate peptide agonists from non-agonists. The ipTM is a metric that assesses the accuracy of the interface between proteins in the predicted multimeric complex and is given as a single number per multimer, ranging from 0 (low accuracy) to 1 (high accuracy)^31^. The pLDDT is a measure of confidence output as a per-residue score ranging from 0 (low confidence) to 100 (high confidence). Since the transmembrane regions of GPCRs are typically predicted with high confidence by AF2, we focused on confidence metrics of the peptide. pLDDT was averaged across peptide residues to yield a single aggregated “peptide pLDDT” score for each GPCR-peptide multimer. Lastly, the PAE quantifies the expected positional error of each residue in a pairwise alignment with every other residue, output as a 2D array with values ranging from 0 Å (low positional error) to a cap at 31.75 Å (high positional error). Per-residue PAE scores for the peptide aligned on the GPCR binding pocket residues were averaged to yield the “pocket PAE” score. For each GPCR-peptide complex, we selected the model from the five AF-Multimer outputs with the highest ipTM, highest peptide pLDDT, or lowest pocket PAE, depending on the specific metric being analyzed.

We evaluated the predictive capability of each metric using mean average precision (mAP). For each GPCR, average precision (AP) measures how agonists rank relative to non-agonists. mAP is then calculated by averaging these AP scores across all receptors, ensuring that GPCRs with few agonists contribute equally to the metric. This approach avoids biases inherent in methods that assign disproportionately higher weight to receptors with many agonists and reduces sensitivity to class imbalance inherent in the dataset. Furthermore, by prioritizing agonists at the top of the ranked peptide list, AP better reflects practical screening scenarios. Analysis of the AP for AF-Multimer confidence metrics (peptide pLDDT, ipTM, and pocket PAE) revealed their ability to discriminate agonists from non-agonists (**Figure 1F, Supplemental Figure 1C-D**). Peptide pLDDT significantly outperformed ipTM and pocket PAE, establishing it as the most reliable indicator of GPCR-peptide agonist interactions among the tested metrics.

### Active-state GPCR-peptide complexes enhance the ability of peptide pLDDT to discriminate agonists

AF2 is inherently biased towards modelling GPCRs in an inactive conformation^34^, likely due to having been exposed to predominantly inactive-state GPCR structures during training. In contrast, agonist binding stabilizes the active conformation of GPCRs^37^. Therefore, we hypothesized that biasing the GPCR in each GPCR-peptide complex toward an active conformation would preferentially improve the confidence of AF-Multimer predictions for agonists.

To model active-state GPCR-peptide complexes, we generated predicted active-state GPCR structures using AF-Multistate^34^ and refined them for use as templates (**Figure 2A**). First, we trimmed extracellular regions from the template, as these domains mediate ligand binding^35^. This allows AF-Multimer to model peptide interactions without interference from pre-defined extracellular structures. Second, we preserved the transmembrane helices, as conformational shifts to the active state are driven by their displacement^38–40^. Third, we removed low-confidence residues in the intracellular domain (involved in G-protein interactions), preventing these regions from distorting structural predictions of active-state GPCR-peptide complexes.

**Figure 2:**
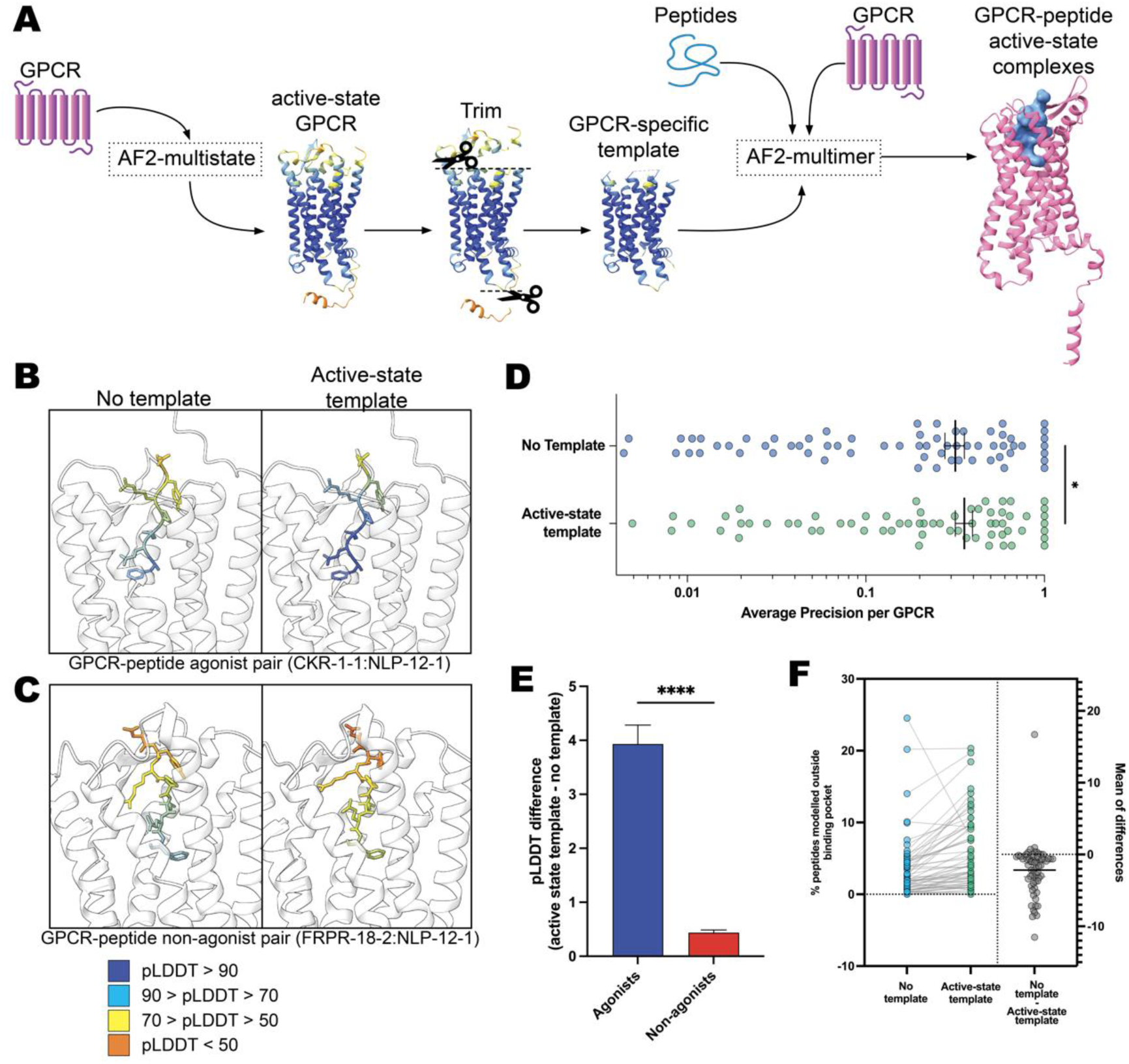
Active-state templates enhance AF-Multimer’s discrimination of agonist and non-agonist GPCR-peptide interactions. (**A**) Predicted active-state structures were generated for each GPCR using AF-Multistate. After trimming all extracellular regions and low confidence (pLDDT < 70) intracellular regions, these templates were then used in AF-Multimer to model GPCR-peptide complexes. (**B**) Representative agonist complex: GPCR CKR-1-1 is modelled with agonist peptide NLP-12-1 without (left) and with (right) an active-state template. The use of an active-state template improved peptide pLDDT. (**C**) Representative non-agonist complex: GPCR FRPR-18-2 is modelled with non-agonist NLP-12-1 (the same peptide shown in B) with (left) and without (right) an active-state template. The use of an active-state template does not alter peptide pLDDT. (**D**) Average precision (AP) of agonist peptides ranked by peptide pLDDT significantly improved when using active-state templates (mAP: 0.36 vs. 0.32 without templates; Wilcoxon signed-rank test, p < 0.05). Error bars indicate mean ± SEM. (**E**) The improvement in peptide pLDDT with active state templates was significantly greater for agonists compared to non-agonists (Welch’s t-test, p < 0.0001). Error bars represent mean ± SEM. (**F**) Active-state templates increased the percentage of peptides modelled outside the GPCR binding pocket (mean: 5.5% ±0.6% SEM vs. 3.4% ±0.6% without templates; paired t-test, p < 0.0001).

Active-state templates enhance discrimination of agonist and non-agonist GPCR-peptide interactions. **Figure 2B** illustrates an example where modelling a GPCR-peptide agonist complex with an active-state template increases peptide pLDDT. In contrast, **Figure 2C** shows the same peptide modelled with a GPCR for which it is not an agonist. Here, the active-state template does not alter peptide pLDDT. These differences in peptide pLDDT between agonist and non-agonist complexes improve the ranking of agonists. Active-state templates increase mAP values compared to template-free modelling (**Figure 2D**) by increasing peptide pLDDT for agonists relative to non-agonists (**Figure 2E**). Additionally, AF-Multimer positioned a greater proportion of non-agonist peptides outside of the binding pocket when using active-state templates (**Figure 2F**). In contrast, using active-state templates did not alter the number of agonist peptides modelled outside of the binding pocket (data not shown). These findings—improved agonist peptide pLDDT and increased exclusion of non-agonist peptides from the binding pocket—demonstrate that active-state GPCR templates improve discrimination between agonist and non-agonist complexes. Consequently, we employed active-state templates for all subsequent analyses.

### Pair representations outperform single representations in discriminating GPCR-peptide agonists

In the process of predicting the 3D structure of proteins, AF2 generates “single” and “pair” representations that are transformed and updated in the EvoFormer block of the algorithm^26^. Each single representation is a matrix in ℝ^{*n* × 256}^ where *n* is the number of residues in the GPCR-peptide complex; similarly, each pair representation is a tensor in ℝ^{*n* × *n* × 128}^, providing pairwise residue-level relationships (**Figure 3A**). These representations can be collected from AF2 as the final hidden layer embeddings of its neural network.

**Figure 3:**
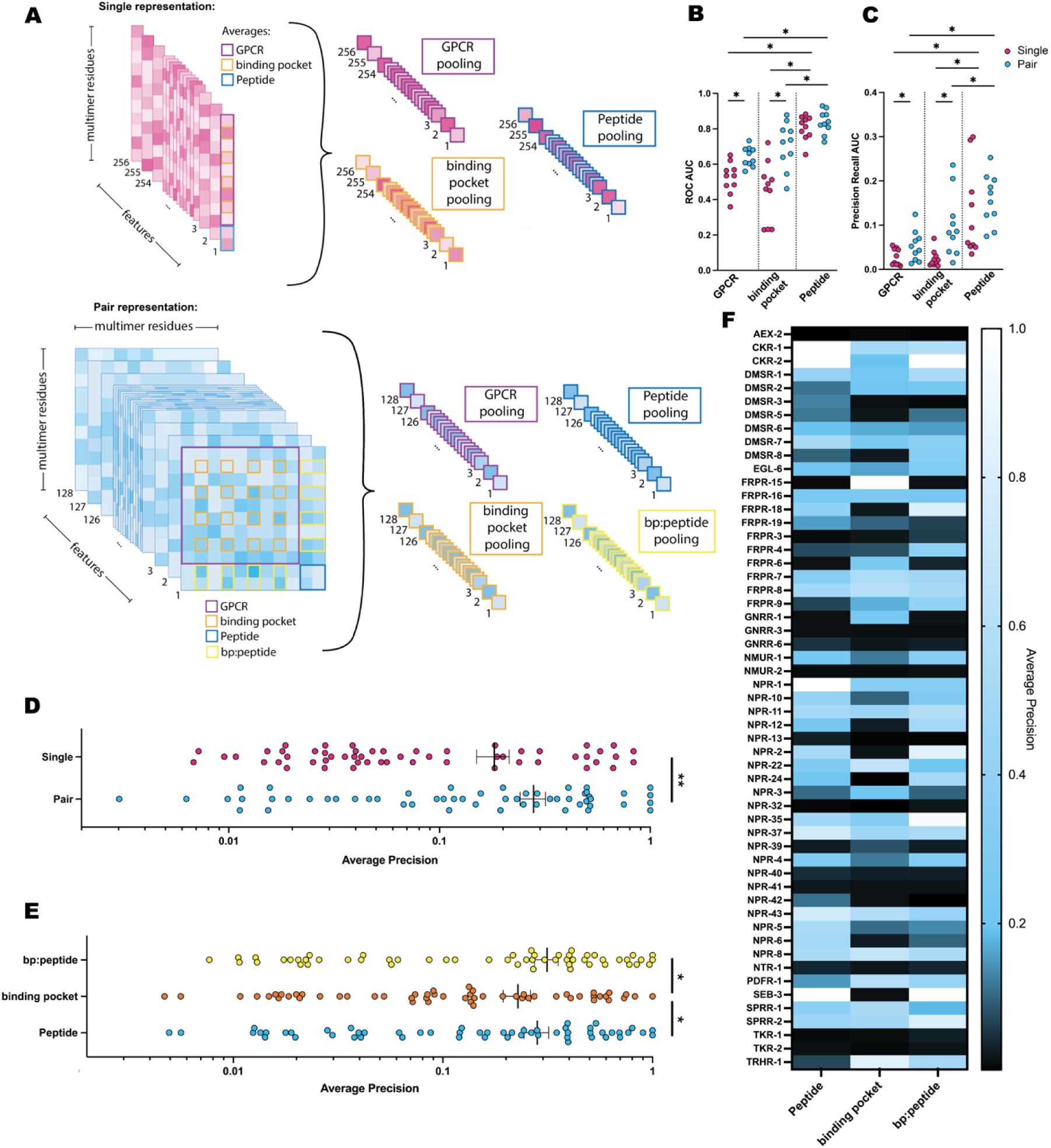
Subregions of AF-Multimer pair representations provide complementary information for classifying agonist and non-agonist peptides. (**A**) Schematic illustration of feature extraction from AF-Multimer representations. Single representations (ℝ^{*n* × 128}^) and pair representations (ℝ^{*n* × *n* × 128}^) were processed by aggregating features via local pooling over predefined subregions: peptide residues (blue outlines), GPCR residues (purple outlines), binding pocket residues (orange outlines), and the pairwise interaction of binding pocket and peptide residues (“bp:peptide”; yellow outlines; pair representations only). (**B**) Pair representations significantly outperform single representations for GPCR and binding pocket subregions, and peptide subregions outperform GPCR and binding pocket subregions. Random forest models were evaluated by ROC-AUC in 10-fold cross validation. Two-way repeated measures ANOVA with Geisser-Greenhouse correction shows significant interaction between the representation type and the subregion used F (1.387, 12.49) = 10.22, p = 0.0044). Multiple comparisons comparing each group to every other group corrected by the FDR method of Benjamini, Krieger, and Yekutieli reveals that peptide subregion performance exceeded that of GPCR or binding pocket subregions, but did not show a significant difference between single and pair representations (* q < 0.05). (**C**) PR-AUC analysis mirrors ROC-AUC results. A two-way repeated measures ANOVA with Geisser-Greenhouse correction was conducted to investigate the effects of representation type and subregion on PR-AUC. A significant main effect was found for representation type (F (1.000, 9.000) = 13.76, p = 0.0048) and subregion (F (1.408, 12.67) = 15.91, p = 0.0008). The interaction between representation type and subregion was not significant (F (1.889, 17.00) = 1.758, p = 0.203). Multiple comparisons comparing each group to every other group corrected by the FDR method of Benjamini, Krieger, and Yekutieli reveals that peptide subregion performance exceeded that of GPCR or binding pocket subregions, but did not show a significant difference between single and pair representations (* q < 0.05). (**D**) A more in-depth evaluation using leave-one-group-out cross-validation (GPCR gene families as groups) reveals that random forest models trained on peptide pair representations significantly outperform single representations (** p < 0.01, Wilcoxon matched-pairs signed rank test). The average precision (AP) was calculated for each GPCR gene. Error bars denote mean ± SEM. (**E**) Leave-one-group-out analysis revealed that random forest models trained on peptide and bp:peptide regions outperformed models trained on the binding pocket subregion (* q < 0.05, Friedman test with multiple comparisons corrected by the FDR method of Benjamini, Krieger, and Yekutieli). (**F**) Heatmap of AP values from (E) illustrates that different GPCR gene families benefit from features provided by different subregions.

We applied local pooling to residue subregions in each representation to identify features useful for classifying GPCR-peptide agonists. For single representations, we averaged residue regions corresponding to the GPCR, the binding pocket, or the peptide, producing three different 256-dimensional feature vectors. Similarly, for pair representations, we averaged residue embeddings involving the same regions and additionally the pairwise intersection of the binding pocket and peptide residues (“bp:peptide”), yielding four distinct 128-dimensional feature vectors.

To evaluate the relative importance of features across subregions of the single and pair representations, we employed random forest classifiers trained using stratified group 10-fold cross-validation. Grouping by GPCR gene ensures that GPCRs are not split across training/validation folds, thereby mitigating data leakage and preventing overfitting to GPCR-specific structural patterns. Additionally, stratifying by agonist class helps reduce class imbalance. Our analysis demonstrates that features derived from the peptide subregions (in both single and pair representations) outperform those from GPCR and binding pocket subregions as measured by receiver operating characteristic (ROC) (**Figure 3B**) and precision-recall (PR) (**Figure 3C**) area under the curve (AUC). While pair representations averaged over GPCR residues or binding pockets significantly outperformed single representations from the same regions in ROC-AUC and PR-AUC, peptide pair and single representations showed no significant difference in these metrics. However, our dataset exhibits significant variation in agonist distribution: some promiscuous GPCRs have many agonists, while most receptors have few (**Figure 1B**). Metrics like ROC-AUC and PR-AUC weight each GPCR-agonist complex equally, which disproportionately prioritizes promiscuous GPCRs in evaluation. To address this, we used the recommender system metric AP, which is computed per GPCR gene such that each gene contributes equally regardless of agonist count or the number of GPCR isoforms derived from that gene. Using leave-one-group-out analysis, we show that peptide pair representations achieve higher AP scores than single representations (**Figure 3D**).

### Distinct subregions of pair representations capture complementary information relevant to successful classification of GPCR-peptide agonists

Our analysis of different subregions in single and pair representations revealed that pair representations provide valuable information for predicting GPCR-peptide agonists. To further investigate this, we trained leave-one-group-out random forest models on three key subregions of the pair representations: (1) the bp:peptide subregion (pairwise residues between the binding pocket and the peptide), (2) the peptide subregion, and (3) the binding pocket subregion.

Models trained on the binding pocket subregion exhibited significantly lower AP scores compared to models trained on the bp:peptide or peptide subregions (**Figure 3E**). However, despite these lower scores, binding pocket-trained models demonstrated unique prioritization capabilities for certain agonists that eluded models trained on other subregions (**Figure 3F**). Notable examples include the GPCRs FRPR-15, GNRR-3, and NPR-3, where binding pocket features proved particularly informative. Similarly, peptide-trained models showed superior performance for some GPCRs, while bp:peptide-trained models were more effective for others. This pattern of variable performance across subregions suggests that features from different subregions are complementary rather than universally optimal for all GPCRs.

### Integration of multimodal GPCR-peptide features for prioritizing agonist interactions

Given the complementary nature of features from different subregions, we generated graph representations of GPCR-peptide complexes (**Figure 4A**) to leverage the ability of GNNs to model complex relational data (e.g., residue interactions).

**Figure 4:**
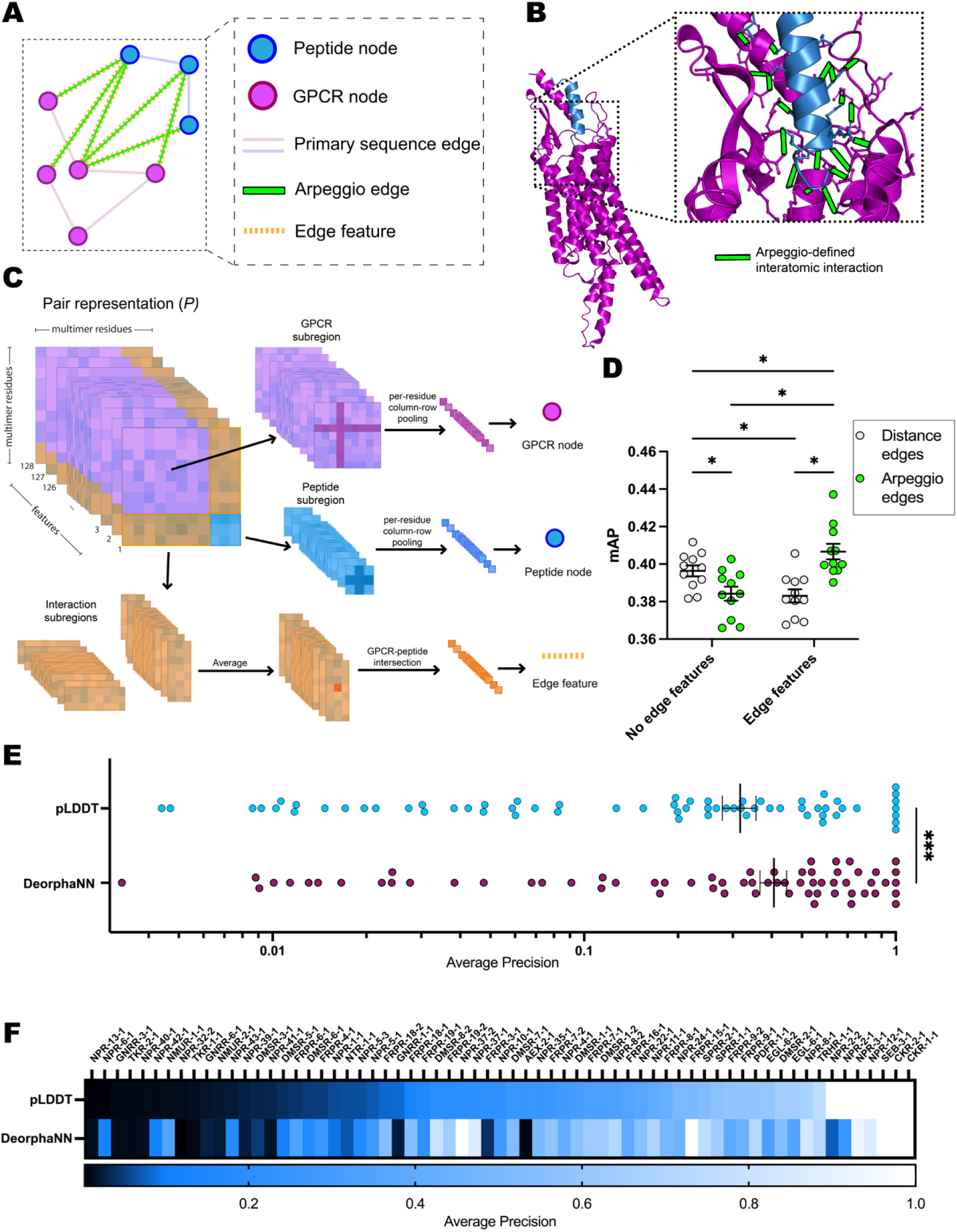
DeorphaNN integrates predicted active state complexes, interatomic interactions, and protein representations to prioritize GPCR-peptide agonists. (**A**) Graph representations of GPCR-peptide complexes. Intra-protein edges are defined by primary sequence, while inter-protein edges are defined by Arpeggio. Peptide residues (all included) and GPCR residues (those directly connected to peptides via Arpeggio-identified interactions or within one degree of such interactions) serve as nodes. (**B**) AF-Multimer predicted structures are analyzed by Arpeggio, which identifies biophysically meaningful interatomic interactions (e.g., hydrogen bonds, ionic interactions). Arpeggio interactions are used to define intermolecular edges between GPCR and peptide residue nodes. (**C**) AF-Multimer pair representations are split into GPCR, peptide, and interaction subregions. Residue-specific pooling of GPCR and peptide subregions generates 128-dimensional embeddings for respective nodes. Interaction edges are annotated with pairwise interaction embeddings from the interaction subregion. (**D**) Arpeggio-defined edges and use of interaction edge features synergistically improve model performance as determined by shuffle -groups-out cross-validation with 11 splits. Two-way repeated measures ANOVA with Geisser-Greenhouse correction shows significant interaction between the use of edge features (no features vs. pairwise interaction embeddings), and method of drawing edges (distance -based cutoff vs. Arpeggio) (F(1, 10) = 28.75, p = 0.0003). Multiple comparisons comparing all groups, corrected by the FDR method of Benjamini, Krieger, and Yekutieli, reveal that loss of edge features and/or Arpeggio defined edges significantly reduces model performance (* q < 0.05). Error bars indicate mean ± SEM. (**E**) DeorphaNN outperforms peptide pLDDT in agonist prediction (Wilcoxon matched-pairs signed-rank test, *** p < 0.001). Error bars indicate mean ± SEM. (**F**) Heatmap of AP values demonstrates that DeorphaNN identifies agonists missed by peptide pLDDT across various GPCR families

The graph representations consist of two distinct sets of nodes: GPCR residue nodes and peptide residue nodes. To define intermolecular edges between these nodes, we use Arpeggio^41^, a computational tool designed to identify biophysically meaningful interatomic interactions in protein structures by analysing spatial and physicochemical relationships (e.g., hydrogen bonds, ionic interactions, cation-π interactions, etc.) (**Figure 4B**). Only GPCR residues directly connected to peptide residues via Arpeggio-identified interactions (or within one degree of such interactions) are selected as nodes, ensuring the graphs focus on biophysically relevant regions of each GPCR. In contrast, all peptide residues are included as nodes, as the smaller size of peptides enables their inclusion without excessive computational burden. Additionally, intramolecular connectivity is preserved by connecting nodes with edges based on their primary sequences.

GPCR nodes are loaded with embeddings derived by residue-specific pooling of the pair representation’s GPCR subregion, while peptide nodes are loaded with embeddings generated similarly from the peptide subregion (**Figure 4C**). Specifically, for each residue *i*, we average the *i-*th row and th*e i*-th column across all features of the respective subregion matrix, yielding a 128-dimensional embedding that captures residue-level structural and interaction information from the pair representation. Edges between GPCR and peptide residues are loaded with pairwise interaction embeddings. This design ensures that node embeddings encode residue-specific information, incorporating both local and global context from the pair representation, while interaction edges encode intermolecular residue-residue interaction context.

### Impact of graph structure and edge embeddings on GNN performance

To assess how graph structure and edge features influence performance, we conducted an ablation study (**Figure 4D**). First, we replaced Arpeggio-defined interaction edges with distance-based edges (residues ≤ 6 Å apart), resulting in a significant reduction in AP when evaluated via shuffle-groups-out cross-validation (11 splits). Next, we removed pairwise interaction embeddings from Arpeggio-defined edges, which similarly decreased performance (**Figure 4D**). Interestingly, when both changes were combined—using distance-based edges and omitting edge features—the model outperformed either single ablation alone, although this combined ablation still achieved significantly lower AP than the complete graph structure (Arpeggio edges with edge features).

Distance-based edges, while capturing spatial proximity, may introduce non-functional contacts that create noise when paired with pairwise interaction embeddings, likely because they lack the physicochemical specificity inherent in Arpeggio-defined interactions. Conversely, while Arpeggio edges provide biologically relevant connections, the model cannot effectively make use of this connectivity without the context provided by pairwise interaction embeddings. The complete graph structure, which combines Arpeggio-defined edges with edge features, outperformed all ablated versions, suggesting a synergistic effect of biophysically meaningful edges and interaction-aware features. This framework enables the GNN, DeorphaNN, to learn complex relational patterns that achieve significantly higher AP scores than a peptide pLDDT-based approach (**Figure 4E**). Importantly, DeorphaNN can prioritize agonists that are overlooked by peptide pLDDT (**Figure 4F**), demonstrating its ability to identify agonists independently of pLDDT-based prioritization.

### Validation on phylogenetically diverse data and comparison to state-of-the-art

To assess whether DeorphaNN captures conserved mechanisms of GPCR-peptide agonism rather than species-specific patterns, we evaluated its performance on a dataset from the bilaterian marine annelid, *Platynereis dumerilii*^11^. This dataset contains 18 GPCRs and 122 peptides that were tested in a similar experimental pipeline as the *C. elegans* training dataset (high-throughput CHO Gα_16_-mediated calcium signalling assay). We utilized DeorphaNN to rank *Platynereis* peptides for each GPCR based on predicted agonist potential. The mAP across all *Platynereis* GPCRs was significantly greater than the baseline expected with random ranking (**Figure 5A**), demonstrating that DeorphaNN generalizes to non-*C. elegans* datasets.

**Figure 5:**
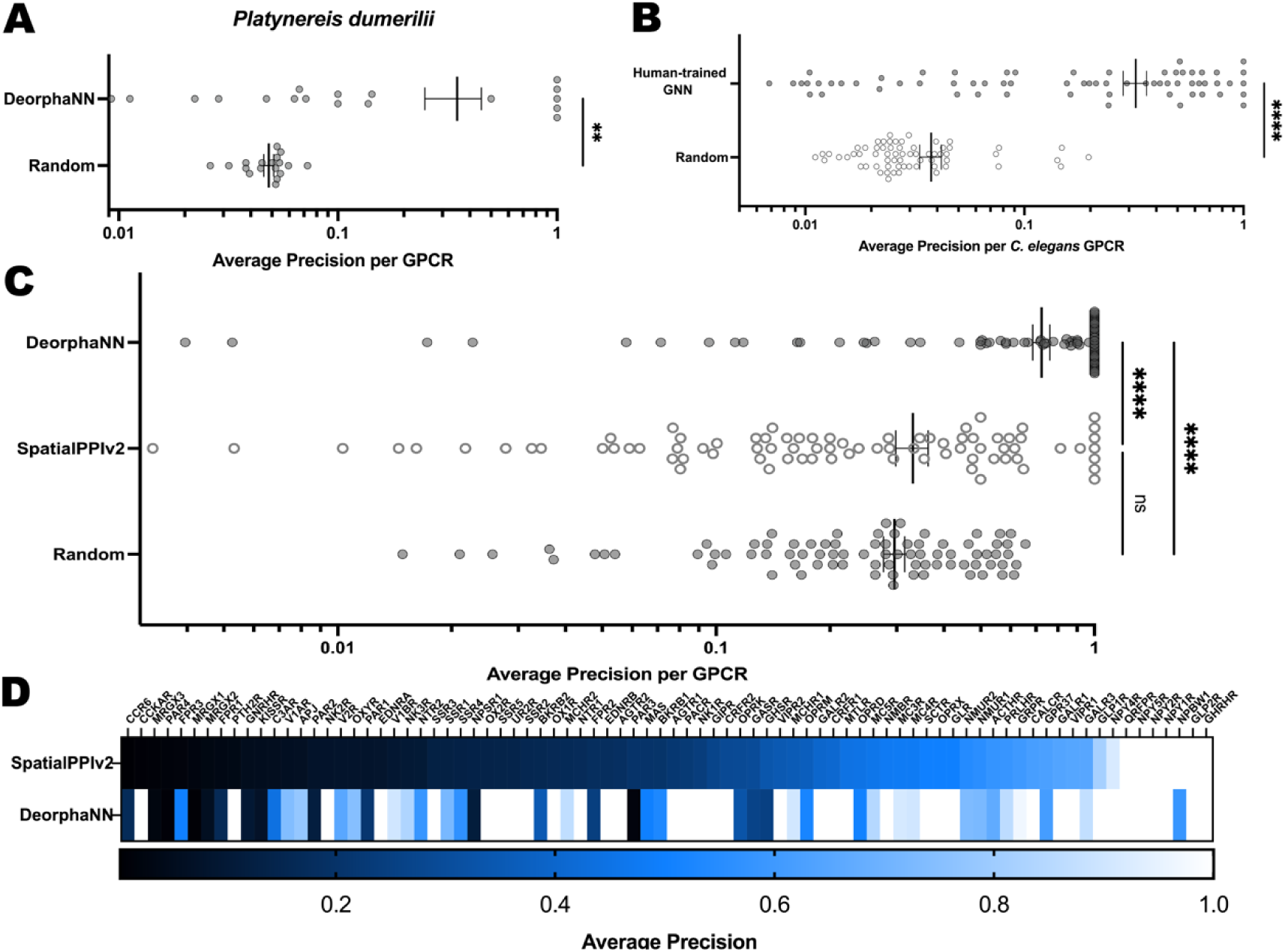
DeorphaNN generalizes to non-*C. elegans* datasets. (**A**) DeorphaNN, trained on *C. elegans* data, prioritizes peptide agonists for GPCRs from *Platynereis dumerilii* significantly better than random ranking. (Wilcoxon matched-pairs signed rank test, ** p < 0.01). (**B**) A human GPCR-peptide agonist dataset was augmented with ESM-2-guided synthetic non-agonist pairs. Its representativeness was evaluated by training a DeorphaNN model on this augmented dataset and assessing its performance on the *C. elegans* dataset, which contains no synthetic data. Model performance was significantly greater than random ranking (Wilcoxon matched-pairs signed-rank test, **** p < 0.0001). (**C**) DeorphaNN (trained on the *C. elegans* dataset) significantly outperforms SpatialPPIv2 and random ranking on human GPCR-peptide agonist prioritization. Statistical significance was determined using the Friedman test with Dunn’s correction (**** p < 0.0001). Error bars indicate mean ± SEM. (**D**) Heatmap for the per-GPCR AP values from (C), illustrating that DeorphaNN prioritizes agonists overlooked by SpatialPPIv2.

To assess DeorphaNN’s generalization to human GPCR-peptide interactions, we utilized a literature-curated dataset of experimentally confirmed agonist interactions across species^42^. While this dataset includes human GPCR-agonist pairs, it lacks non-agonist interactions. To address this gap, we generated synthetic non-agonist GPCR-peptide pairs using a dissimilarity-based approach inspired by principles from prior work^42^. Specifically, we leveraged ESM-2^43^ language model embeddings to quantify peptide dissimilarity. By calculating the Euclidean distance between each peptide’s ESM-2 representation and that of the cognate agonists, we selected peptides with a distance exceeding an empirical threshold—chosen to ensure at least 3 non-agonists per GPCR.

To validate the biological relevance of these synthetic pairs, we trained a model using the DeorphaNN framework on the human dataset (augmented with synthetic non-agonist interactions) and tested its performance on the *C. elegans* dataset, which contained no synthetic data (**Figure 5B**). This model achieved scores significantly higher than the random baseline, confirming that the human dataset—which combines confirmed agonist pairs with ESM-2-guided synthetic non-agonists—reliably captures real-world interaction patterns and supports the augmented dataset’s utility for benchmarking.

We then benchmarked DeorphaNN (trained on *C. elegans* data) on the augmented human dataset, achieving a mAP of 0.72—significantly higher than the random baseline of 0.30 (**Figure 5C**). For comparison, we evaluated SpatialPPIv2, a state-of-the-art model designed for protein-protein interaction prediction that leverages large language models and graph attention networks to capture structural patterns^44^. On the same augmented human dataset, SpatialPPIv2 achieved a mAP of 0.33, not significantly different from random ranking. DeorphaNN significantly outperformed SpatialPPIv2, demonstrating greater accuracy in predicting GPCR-peptide agonist interactions. This difference highlights that our specialized GNN—which integrates structural predictions and interaction data—captures biologically relevant features specific to GPCR-peptide agonism more effectively than general protein-protein interaction models.

### Discovery of novel peptide agonists for orphan GPCRs identified by virtual screening

Some peptide agonists were identified after the system-wide experimental screen of GPCR-peptide interactions in *C. elegans*. Indeed, peptidomics and comparative genomics studies have since expanded the number of peptides known in *C. elegans* beyond those included in the training dataset^45–47^, providing an opportunity to reassess *C. elegans* GPCRs with a broader set of candidate ligands. The receptor NPR-34, an ortholog of the elevenin peptide receptor family, was not found to have agonist-induced activity with any of the peptides in the training dataset, but subsequent experiments demonstrated that this receptor is activated by a *C. elegans* peptide (SNET-1-1) related to the elevenin peptide family^9^. This agonist ranks first out of 364 candidates (339 peptides from the training dataset plus 25 recently identified peptides [**Supplemental Data 1**]) by DeorphaNN’s predictions (**Table 1**). Similarly, the peptide NLP-73-3 was present in the training dataset but did not activate SEB-2 in the original screen because the synthetic peptide lacked a post-translational modification (disulfide bridge) important for receptor activation; however, when synthesized with this modification, NLP-73-3 robustly activated SEB-2^9^. Indeed, NLP-73-3 is ranked first among all candidate peptides for SEB-2 by DeorphaNN (**Table 1**).

**Table 1:**
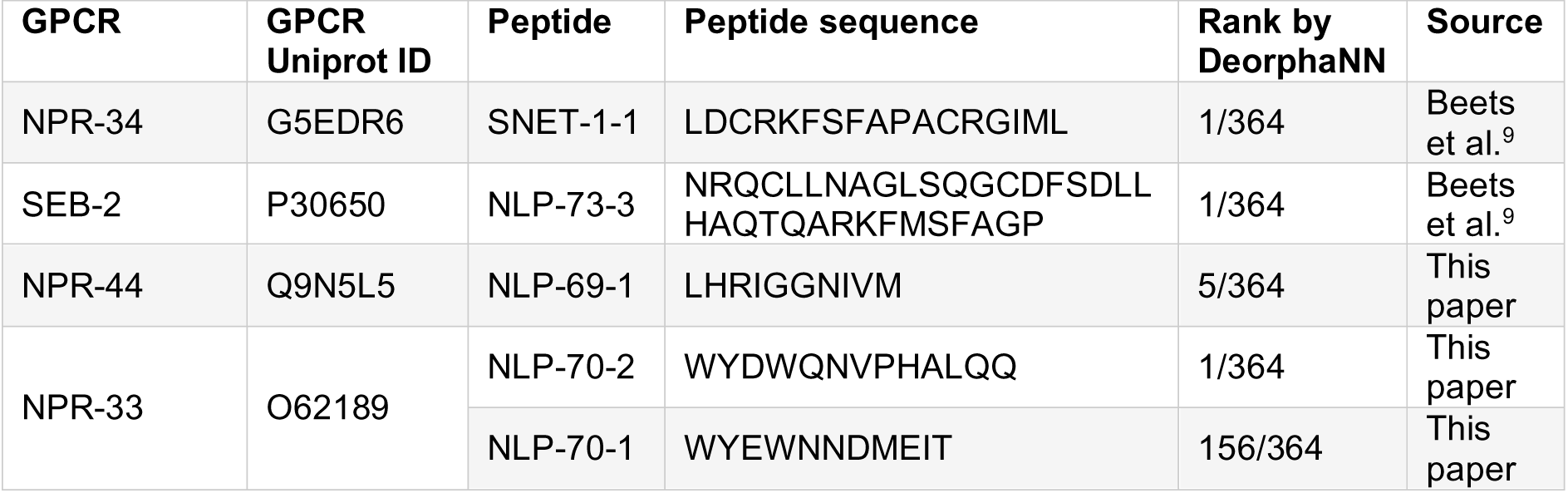
GPCR-peptide agonist pairs identified by DeorphaNN.

We next applied DeorphaNN to two orphan *C. elegans* GPCRs, NPR-44/H23L24.4 and NPR-33/F31B9.1, and tested their highest-ranked candidate agonists. For NPR-44, we observed robust activation by NLP-69-1 (EC_50_ value of 231.7 nM), ranked 5^th^ out of 364 peptides by DeorphaNN (**Table 1**, **Figure 6A, Supplemental Data 3**). In addition, NPR-33 was potently activated by NLP-70-2 (EC_50_ value of 76.3 nM), the highest-ranked peptide (**Table 1**, **Figure 6B, Supplemental Data 3**). No response was seen with any of the peptides in cells transfected with an empty vector control (**Supplemental Figure 2A**). Given that GPCRs are often activated by multiple peptides within the same peptide-encoding gene (**Supplemental Figure 1A**), we also tested NLP-70-1 (ranked 156/364 by DeorphaNN), another putative peptide encoded by the *nlp-70* gene. Although we detected receptor activation, NLP-70-1’s EC_50_ value (1.16 µM; **Figure 6C**) is considerably higher than that of NLP-70-2. This suggests DeorphaNN may prioritize agonists with specific interaction patterns that correlate with higher potency. Indeed, the majority of agonist interactions in the *C. elegans* dataset exhibit high potency (EC_50_ < 500 nM)^9^, which may explain why lower-potency agonists like NLP-70-1 were ranked lower by DeorphaNN.

**Figure 6:**
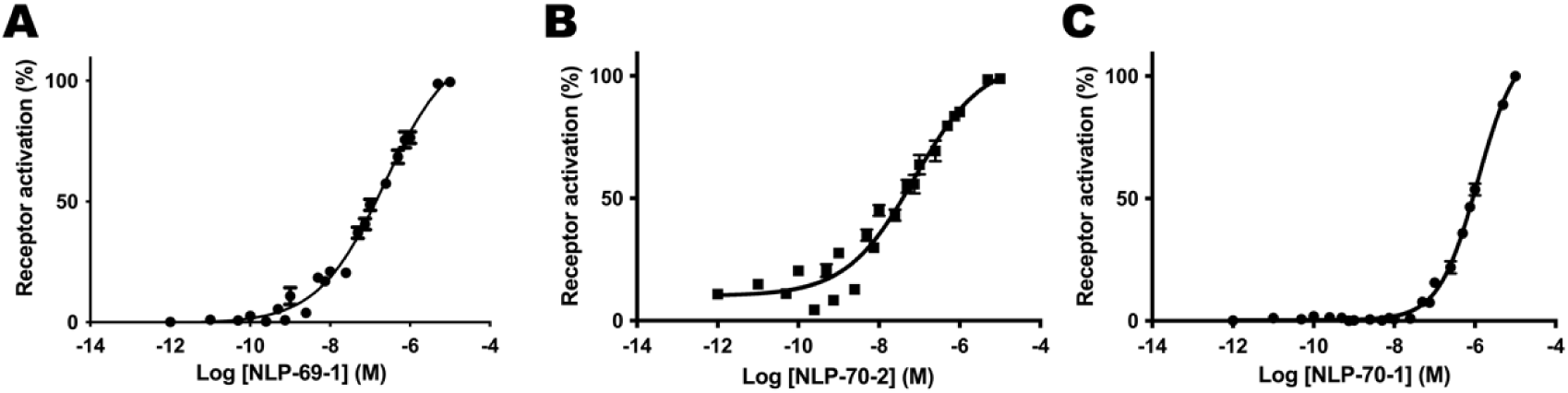
DeorphaNN-guided identification and experimental validation of agonists for two *C. elegans* GPCRs. (**A**) Concentration-response curve showing calcium mobilization in CHO cells co-expressing the GPCR NPR-44, Gα16, and the calcium indicator aequorin upon stimulation with the peptide NLP-69-1. Responses are shown relative to 100% activation after normalization to the total calcium response. EC₅₀ = 231.7 nM (95% CI: 153.6 nM - 309.7 nM), n = 6. (**B**) Concentration-response curve of the GPCR NPR-33 to the peptide, NLP-70-2. EC₅₀ = 76.3 nM (95% CI: 50.3 nM - 128 nM), n = 6. (**C**) Concentration-response curve of the GPCR NPR-33 to the peptide NLP-70-1. EC₅₀ = 1.16 µM (95% CI: 1.00 µM - 1.38 µM), n = 6.

## DISCUSSION

In this study, we present DeorphaNN, a GNN-based pipeline for predicting GPCR-peptide agonist interactions *in silico* that offers a scalable strategy for GPCR deorphanization. Previous studies have shown that pLDDT from AlphaFold predictions correlate with interaction affinity^31,32,48^, consistent with our findings that pLDDT partially discriminate agonists from non-agonists. However, affinity alone is insufficient to determine agonist function^35,49^. While affinity is a prerequisite for agonism, it does not capture additional mechanisms required for functional activation, such as intrinsic efficacy^50^—the propensity of a ligand to induce conformational changes in the GPCR that activate downstream G protein signalling.

Agonist binding induces conformational changes in the GPCR that drive conserved transmembrane (TM) rearrangements, a hallmark of receptor activation^38,51,52^. A key feature of this process is the outward movement of TM6^53^, facilitated by kinking at conserved motifs and disruption of ionic interactions between TM3 and TM6/7^54–56^. These rearrangements reconfigure the surface of the orthosteric binding pocket^57,58^, enhancing affinity for agonists^59^. To capitalize on this mechanism, DeorphaNN incorporates active-state structural templates generated using AF-Multistate^34^. This enables the modelling of GPCR-peptide complexes in their active conformations, leveraging AF-Multimer’s ability to prioritize peptides that both bind and stabilize the active-state binding pocket.

DeorphaNN builds on AF-Multimer by analyzing patterns in the predicted active-state complex to predict whether bound peptides can activate receptors. Agonists activate GPCRs through interactions with key “toggle switch” residues in the binding pocket^60,61^. This activation is thought to require two distinct types of intermolecular interactions: “anchor” interactions that provide stability for ligand binding, and “driver” interactions that directly engage with toggle switches to induce GPCR conformational changes^62–64^. To enable pattern recognition of these interactions, we engineered DeorphaNN to utilize Arpeggio-defined edges—capturing interatomic interactions^41^—and AF pairwise embeddings— encoding implicit biophysical relationships between residues^27^. By integrating these features, DeorphaNN learns to distinguish agonists from non-agonists, likely through patterns corresponding to the anchor and driver interactions essential for receptor activation and intrinsic efficacy. In contrast, SpatialPPIv2—a state-of-the-art model for predicting general protein-protein interactions^44^—lacks this context, explaining its inferior performance in differentiating agonist from non-agonist interactions (**Figure 5C-D**).

Despite being trained on a relatively small number of GPCR-agonist pairs from *C. elegans*, DeorphaNN’s predictions seem to generalize across species. This success reflects the effective application of inductive transfer learning principles, which have revolutionized fields like natural language processing and computer vision by enabling models pretrained on large datasets to be fine-tuned for specialized tasks with limited data^65,66^. In our work, DeorphaNN leverages the AF2 algorithm, which was pretrained on approximately 170,000 protein structures from diverse species^26^, and adapts it for the specialized task of GPCR-peptide agonism prediction. The model’s ability to predict agonist interactions in phylogenetically distinct species such as *Platynereis dumerilii* and humans suggests that the implicit biophysical principles learned by AF2 were successfully transferred to this specialized task.

Compared to existing methods, DeorphaNN represents a significant advancement in GPCR-peptide agonism prediction. PD-incorporated SVM, a sequence-based approach, predicts peptide agonists for GPCRs using a support vector machine trained on peptide descriptors that represent sequence patters of 1-5 residues encoded into numeric arrays^42^. GPCRVS employs deep neural networks, gradient boosting machines, and AutoDock Vina to predict compound activity, selectivity, and binding affinity for small molecules and peptides, but is restricted to GPCRs within its training data and can only effectively screen N-terminal fragments of peptides (six residues or smaller)^67^. In contrast, DeorphaNN integrates structural predictions from AF-Multimer, deep learning-derived protein representations, and intermolecular interactions data, enabling it to learn from three-dimensional structural relationships and biophysical interaction patterns that are not captured by primary sequence alone. Trained on biologically relevant peptide sizes (3-37 residues) and demonstrating generalization to non-*C. elegans* GPCRs, DeorphaNN provides a more comprehensive approach to GPCR-peptide agonism prediction across diverse receptors.

Despite its advances, DeorphaNN does not uncover all GPCR-peptide agonist interactions. Several factors likely contribute to this limitation. First, as the training dataset was derived from a single experimental screen, the accuracy of the dataset is inherently tied to the limitations of that screen. For example, while the promiscuous Gα_16_ subunit enables many GPCRs to signal through the PLCβ pathway, the effectiveness of this signalling can be ligand-specific^68^, potentially resulting in misclassified negative interactions. In addition, some peptides may be misclassified for highly promiscuous receptors, as not all peptides from the same precursor gene were tested against these receptors for agonist-induced activity. Second, while the dataset includes a variety of GPCR-peptide agonist interactions, it is not exhaustive, creating potential blind spots for interactions that are dissimilar to those in the training dataset. Third, DeorphaNN’s performance depends on AF2’s ability to accurately predict GPCR-peptide complexes. Inaccurate structural predictions may fail to capture the true nature of GPCR-peptide relationships, impacting DeorphaNN’s effectiveness. Indeed, AF2-predicted GPCR structures show deviations in ligand-binding pockets and extracellular domains compared to experimentally determined structures^69,70^. Additionally, AF2 cannot account for post-translational modifications of GPCRs and peptides. For GPCRs, extracellular post-translational modifications such as tyrosine sulfation, glycosylation, and proteolytic cleavage may alter ligand binding^71^; for peptides, post-translational modifications such as C-terminal amidation or pyroglutamation are often important for agonist activity^35^. Although direct interactions between GPCRs and the C-terminal amidated group of peptide ligands have not yet been identified^35^, amidation has been shown to influence biological activity^72^, and the omission of such post-translational modifications in AF2 predictions may result in lost interaction information. However, with the advent of newer models such as AlphaFold3^73^, Chai-1^74^, and Boltz-1^75^—which can account for post-translational modifications and demonstrate higher modelling accuracy—these limitations may be addressed in future work.

In this study, we demonstrate how deep learning methods can enhance the identification of functional GPCR-ligand relationships, using DeorphaNN to identify agonists for two previously orphan GPCRs in *C. elegans*. Our results demonstrate that DeorphaNN’s predictive capabilities extend beyond this model organism, suggesting its potential to aid in deorphanization efforts of both orphan GPCRs and orphan peptides. Beyond facilitating the discovery of novel signalling pathways and potential therapeutic targets, DeorphaNN may significantly accelerate drug discovery efforts, as recent advances in machine learning-based synthetic peptide design have produced tools for generating synthetic ligands for GPCRs^76,77^. DeorphaNN complements all of these efforts, prioritizing promising candidate peptide agonists for experimental validation and reducing the need for exhaustive screening.

## MATERIALS AND METHODS

### Datasets

The *C. elegans* training dataset was derived from a previously published system-wide screen using a CHO Gα_16_-mediated calcium signalling assay to assess endogenous peptide agonist activity across *C. elegans* GPCRs^9^. To accommodate AF-Multimer’s limitations regarding post-translational modifications, we preprocessed peptide sequences by removing C-terminal glycine residues for amidated peptides and N-terminal glutamine residues for pyroglutamated peptides. Peptides containing disulfide bridges were not changed. After removing duplicate sequences and peptides shorter than 3 residues, 339 unique peptides remained. GPCRs that lacked any concentration-dependent agonist responses in the screen were excluded, reducing the likelihood that an observed lack of agonist activity was the result of experimental limitations. Additionally, we excluded the GPCR DMSR-8-1, which has the same agonist interaction profile as the other DMSR-8 isoform (DMSR-8-2) and only differs in sequence by an addition of four N-terminus residues. This filtering process resulted in a final training dataset comprising 65 unique GPCRs derived from 55 genes. The full list of GPCRs and peptides included in the *C. elegans* training dataset is provided in **Supplemental Data 1**.

The *Platynereis dumerilii* dataset used for cross-species validation was derived from a previously published report^11^. We processed all peptides with modified residues as above and omitted peptides with D-amino acids, resulting in 122 peptides. We similarly omitted GPCRs that had no confirmed agonists among the included peptides, resulting in 18 receptors and 23 agonist interactions.

For the human dataset, we utilized a previously published list of GPCR-peptide agonist interactions compiled across diverse species, including vertebrates and inverterbates ^42^. We limited our analysis to agonist interactions involving human GPCRs, while using peptides from all species (excluding those longer than 50 residues) to augment the dataset with synthetic non-agonist interactions. Specifically, we generated ESM2-150M^43^ embeddings for each peptide and identified, for each GPCR, peptides with Euclidean distances > 3.9 from known agonists, ensuring at least 3 putative non-agonists per GPCR. This process yielded an augmented dataset containing 345 agonist and 3370 non-agonist GPCR-peptide pairs across 82 human GPCRs.

In all datasets, we omitted GPCRs that were not predicted by DeepTMHMM to have the characteristic GPCR structure (seven transmembrane domains, with an extracellular N-terminal and an intracellular C-terminal). The full list of GPCRs and peptides included in the validation datasets (*Platynereis* and human) are provided in **Supplemental Data 2**.

### AlphaFold2

Primary amino acid sequences for each GPCR-peptide complex were input together (GPCR:peptide) into AF-Multimer^26,31^ using local ColabFold v1.5.5^78^ on a high-performance computing cluster. All modelling settings were left at default (model_type=alphafold2_multimer_v3, num_recycles=20, recycle_early_stop_tolerance=0.5, pairing_strategy=greedy, max_msa=auto), resulting in the generation of 5 predicted structures and corresponding confidence metrics. Hidden layer protein representations (single and pair) were collected.

For confidence metric analyses, we utilized the model with the highest ipTM, highest peptide pLDDT, or lowest pocket PAE, depending on the metric being analysed. To determine whether the peptide was modelled within the binding pocket, we carried out analysis on the predicted structure with the highest peptide pLDDT, measuring the minimum distance of peptide to binding pocket residues after averaging the location of all atoms in each residue.

For all protein representation analyses, representations across all five AF-Multimer models were averaged to prevent algorithms from learning patterns specific to the different AF-Multimer models.

To bias AF-Multimer predictions towards GPCR active-state conformations, we first generated active-state GPCR templates using AF2-Multistate^34^. This approach leverages state-annotated GPCR databases to guide predictions with a restricted MSA protocol. The top ranked active-state structures were processed by trimming intracellular residues with pLDDT values below 70 and all extracellular residues, based on DeepTMHMM^36^ annotations. These processed structures served as GPCR-specific templates in subsequent AF-Multimer predictions for each GPCR-peptide pair, generating state-biased predictions. MSA reduction was not necessary as the templates were derived from the same sequence as the target GPCRs^79^, and retaining the full MSA preserves coevolutionary signals that are vital for driving accurate inter-protein contact predictions^80^.

### Mean Average Precision (mAP)

Ranking performance was quantified according to recommendation system metrics^81^, using average precision (AP) computed across all peptides (*N*) per GPCR or GPCR gene. For each receptor, AP aggregates precision at every rank containing an agonist:

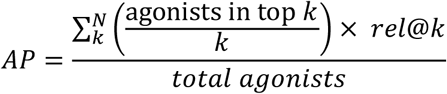

where *rel@k* equals 1 if the *k*-th ranked peptide is relevant (i.e. an agonist) and 0 otherwise. Mean average precision (mAP) averages AP equally across all GPCRs.

To establish a random baseline, we randomized peptide rankings 100 times per GPCR or GPCR family and computed the AP. The average of the 100 APs was then used as the random baseline AP.

### Supervised learning algorithms

We implemented random forest classifiers using scikit-learn^82^ with default hyperparameters, leveraging their ability to perform implicit feature selection and assess feature importance. To maintain consistency across all cross-validation splits, we used identical random seed and data partitions. For Figures 3B and 3C, we employed stratified group 10-fold cross-validation with the same random seed. Grouping by GPCR was performed to minimize leakage of receptor-specific information between folds. Isoforms from the same GPCR gene were treated as a single unit, ensuring they appear together (55 unique GPCR genes), and class distribution was balanced by stratification. For Figures 3D and 3E, we employed leave-one-group-out cross-validation. Reported APs are an average of 5 different random seeds.

### Arpeggio

The top-ranked GPCR-peptide structure according to peptide pLDDT was examined. Peptides that exhibited a minimum distance greater than 12.5 Å from the binding pocket were excluded from further analysis. The position of each residue is defined by averaging the position of each residue atom. Structures were relaxed by Amber^83^ and analysed with Arpeggio^41^ to identify interatomic interactions between the GPCR and peptide. Specifically, we identified hydrophobic interactions, polar bonds, weak polar bonds, hydrogen bonds, weak hydrogen bonds, ionic bonds, aromatic ring interactions, van der Waals’ forces, and amide group interactions.

### Graph construction

Graphs were constructed from GPCR-peptide interaction data obtained from AlphaFold2 structural predictions and pair representations. Each graph represents a GPCR-peptide complex, with nodes corresponding to individual residues. All peptide residues were included as nodes, while GPCR residues were included if they directly interacted with peptide residues or were within one degree of such interactions. Nodes were connected by two types of edges: intramolecular edges connecting nodes according to primary sequence, and intermolecular edges connecting GPCR and peptide nodes according to Arpeggio-identified interactions. For Figure 4D, spatial proximity-based edges (< 6 Å) were also utilized, computed by defining the residue’s position as the average position of the atoms that make up the residue.

Node features were derived using pair representations averaged over 5 models for each residue position. First, we processed the tensor ***P*** ∈ ℝ^*n*×*n*×128^—where *n* is the total residue count in the GPCR-peptide complex—to extract three sub-tensors:

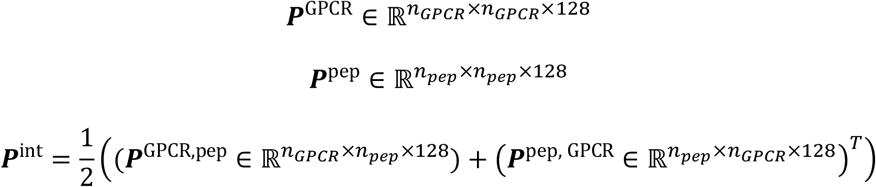

where *n_GPCR_* and *n_pep_* are residues of the GPCR and peptide, respectively.

We derived per-residue embeddings for GPCR and peptide nodes by applying cross-axis pooling independently to sub-tensors, ***P***^GPCR^ and ***P***^pep^. For each residue *i* in the sub-tensor *X* ∈ {GPCR, pep}, we computed its embedding by averaging within each of the 128 features across the *i*-th row and the *i*-th column while correcting for (*i*, *i*) duplication:

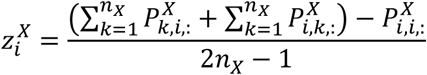

Intermolecular edge features were derived from the interaction region of the pair representations (***P***^int^), with each intermolecular edge loaded with a 128-dimensional feature vector representing the interaction between the residue pairs connected by that edge. Intramolecular edges were assigned a uniform feature vector, computed as the mean-pooled representation of all intermolecular edge features across the graph.

### Model architecture and training

The GNN architecture is based on the Graph Attention Network (GAT) framework, specifically utilizing the GATv2Conv^84^ layer implementation from PyTorch Geometric. The network begins with a Batch Normalization layer to standardize input features, ensuring stable training dynamics. This is followed by a GATv2Conv layer with 10 attention heads, a dropout rate of 0.5, and 64 hidden channels. The attention mechanism allows the model to dynamically weigh the importance of neighbouring nodes differently when aggregating information, enabling it to focus on relevant structural and interaction patterns within the graph.

After the convolutional layer, a rectified linear unit (ReLU)^85^ activation function introduces non-linearity to the model. For graph-level representation learning, a global mean pooling operation aggregates node embeddings across the entire graph, producing a fixed-size vector representation regardless of graph size. This pooled representation is then passed through a final linear layer with a dropout rate of 0.5 for regularization before being fed into a softmax function for classification. The model is trained using the AdamW optimizer^86^ with a learning rate of 0.0005 and employs CrossEntropyLoss^87^ with label smoothing to mitigate overfitting and improve generalization. During training, graphs are batched and processed using PyTorch Geometric’s DataLoader, with edge features incorporated where applicable.

For each validation fold, the training set was subsampled to maintain a balance between positive and negative examples. Specifically, for each positive example (agonist GPCR-peptide complex), four negative examples (non-agonist GPCR-peptide complex) were randomly selected, creating a balanced training set with a 1:4 ratio of positive to negative samples. The model selection was based on validation performance using group shuffle-split cross-validation, with early stopping to prevent overfitting. In this approach, the dataset was divided into groups based on GPCR families, and multiple random splits were generated where each split maintained a specified proportion of groups in the training and validation sets. This method ensures that the model is evaluated on previously unseen GPCR families while maintaining biological relevance in the validation process. The final model for each fold was chosen based on the highest validation average precision score achieved during training. Training was performed for 30 epochs, with the model evaluated periodically on the validation set to monitor convergence and generalization performance.

The final predictions reported were generated using an ensemble of 10 GNNs trained via bootstrap aggregating, with each model trained on a distinct 90% bootstrap sample of the training data. The ensemble outputs were aggregated to produce the final prediction scores.

### Aequorin-based GPCR activation assay

An aequorin-based calcium mobilization assay was used to assess GPCR activation following peptide application, as previously described^9^. In brief, reverse pharmacology screening was performed using CHO-K1 cells expressing mitochondrial-targeted apo-aequorin and promiscuous human Gα_16_. CHO cells were transfected with pcDNA3.1 plasmids containing the GPCR of interest and harvested two days after transfection. BSA served as a negative control for peptide-evoked calcium responses; ATP activates an endogenous receptor in CHO cells and was used as a positive control. Maximum calcium responses for normalization were obtained by lysing cells at the end of the assay.

### Statistical analysis

Statistical analysis was performed in GraphPad Prism version 10.2.2 for macOS (GraphPad Software, San Diego, CA). All statistical tests were two-tailed.

## RESOURCE AVAILABILITY

### Lead Contact

Requests for further information and resources should be directed to and will be fulfilled by the lead contact, Larissa Ferguson.

### Materials Availability

This study did not generate new unique reagents.

### Data and Code Availability

All datasets utilized in this study are available as Supplemental Materials and in a Hugging Face repository at https://huggingface.co/datasets/lariferg/DeorphaNN. The DeorphaNN code is available at https://github.com/Zebreu/DeorphaNN.

## ACKNOWLEDGEMENTS

We thank Jake Grimmett, Toby Darling, and Ivan Clayson of LMB Scientific Computing. Additional thanks to members of the Schafer lab for helpful discussions. Z.W. acknowledges support from the Max Perutz Fund as part of the GenerationResearch project. W.R.S. acknowledges support from the Medical Research Council (MRC) core grant MC-A023-5PB91 and the Research Foundation – Flanders (FWO) grant G050825N. I.B. acknowledges support from the KU Leuven Research Council grant C16/19/003, the Research Foundation – Flanders (FWO) grant G036524N, and the Baillet Latour Fund.

## AUTHOR CONTRIBUTIONS

L.F., S.O., T.K.L., W.R.S., and I.B. conceptualized the work; L.F., S.O., and T.K.L developed methodology; L.F., S.O., and T.K.L. performed formal analysis; L.F. and T.K.L. contributed to visualization; S.O., L.F., and C.W. developed software; L.F., S.O., E.V., C.W., Z.W. conducted investigation and experiments; L.F. wrote the original manuscript draft; T.K.L., I.B., S.O., W.R.S., and C.W. reviewed and edited the manuscript; W.R.S., I.B., and L.F. supervised the project. W.R.S. and I.B. provided funding and infrastructure.

## DECLARATION OF GENERATIVE AI AND AI-ASSISTED TECHNOLOGIES IN THE WRITING PROCESS

During the preparation of this work, ChatGPT and KIMI.ai were used to improve readability. After using these tools/services, the authors reviewed and edited the content as needed and take full responsibility for the content of the publication.

## SUPPLEMENTAL INFORMATION

Supplemental Figures 1-2

<u>Supplemental Data 1</u>: An excel file with the names and primary sequences of all GPCRs and peptides in the *C. elegans* training dataset, and a list of the experimentally validated agonist pairs.

Supplemental Data 2: An excel file with the names and primary sequences of all GPCRs and peptides in the validation datasets (*Platynereis* and human), with agonist pairs labelled.

<u>Supplemental Data 3</u>: An excel file with DeorphaNN predictions (ranked) for SEB-2, NPR-34, NPR-44, and NPR-33 with 364 *C. elegans* peptides

