## Supplemental Figures 1-2 for "DeorphaNN: Virtual screening of GPCR peptide agonists using AlphaFold-predicted active-state complexes and deep learning embeddings"

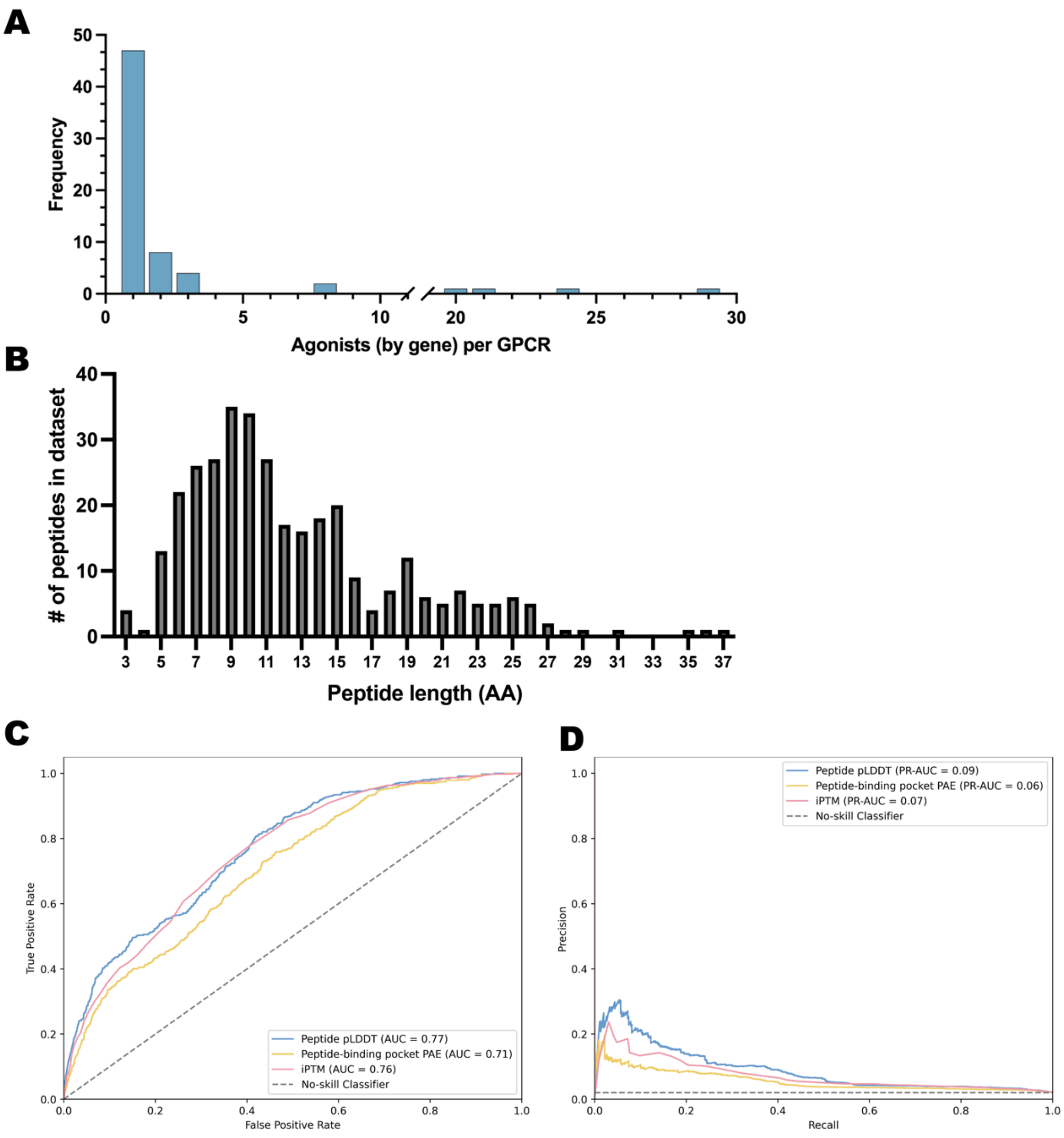


**Supplemental Figure 1**

(**A**) Frequency distribution of the number of agonists by gene per GPCR in the *C. elegans* dataset. (**B**) Frequency distribution of the amino acid count per peptide in the *C. elegans* dataset. (**C**) Receiver operating characteristic (ROC) curves and (**D**) Precision-recall (PR) curves for the ipTM, peptide pLDDT, and pocket PAE confidence metrics, illustrating the ability of these confidence metrics to classify agonist and non-agonist GPCR-peptide pairs from the *C. elegans* dataset.

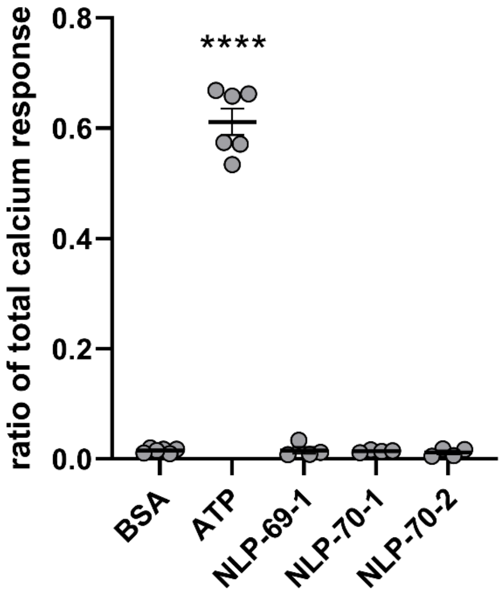


**Supplemental Figure 2**
Peptides do not affect calcium levels in CHO cells in the absence of transfected GPCR. Values are reported as the ratio of the total calcium response in cells transfected with empty vector and challenged with peptides (NLP-70-1, NLP-70-2, or NLP-69-1) (10 µM), BSA (negative control), or ATP (positive control) ($n \geq 4$). Error bars indicate SEM. Significance was assessed by one-way ANOVA with Tukey's multiple comparisons test. No significant difference was observed between BSA and peptide conditions ($p > 0.05$). Calcium response to ATP was significantly higher than BSA or peptides (**** $p < 0.0001$).
